# Real-time tractography-assisted neuronavigation for TMS

**DOI:** 10.1101/2023.03.09.531565

**Authors:** Dogu Baran Aydogan, Victor H. Souza, Renan H. Matsuda, Pantelis Lioumis, Risto J. Ilmoniemi

## Abstract

**Background:** State-of-the-art navigated transcranial magnetic stimulation (nTMS) systems can display the TMS coil position relative to the structural magnetic resonance image (MRI) of the subject’s brain and calculate the induced electric field. However, the local effect of TMS propagates via the white-matter network to different areas of the brain, and currently there is no commercial or research neuronavigation system that can highlight in real time the brain’s structural connections during TMS.

**Objective:** To develop a real-time tractography-assisted TMS neuronavigation system and investigate its feasibility.

**Method:** We propose a modular framework that seamlessly integrates offline (preparatory) analysis of diffusion MRI data with online (real-time) tractography. For tractography and neuronavigation we combine our custom software Trekker and InVesalius, respectively. We evaluate the feasibility of our system by comparing online and offline tractography results in terms of streamline count and their overlap.

**Results:** A real-time tractography-assisted TMS neuronavigation system is developed. Key features include the application of state-of-the-art tractography practices, the ability to tune tractography parameters on the fly, and the display of thousands of new streamlines every few seconds using a novel uncertainty visualization technique. We demonstrate in a video the feasibility and quantitatively show the agreement with offline filtered streamlines.

**Conclusion:** Real-time tractography-assisted TMS neuronavigation is feasible. With our system, it is possible to target specific brain regions based on their structural connectivity, and to aim for the fiber tracts that make up the brain’s networks.

## 1. Introduction

Transcranial magnetic stimulation (TMS) is a non-invasive brain stimulation technique approved in many countries, for treating major depression disorder (MDD) (Fitzgerald et al., 2009), obsessive–compulsive disorder (OCD) (Pelissolo et al., 2016), and for performing motor and speech cortical mapping as part of presurgical evaluation (Lefaucheur and Picht, 2016; Krieg et al., 2017). The navigated TMS (nTMS) approach is also widely used to investigate brain functions by evoking motor or behavioral responses or by interrupting task-related processes (Grosprêtre et al., 2016; Di Lazzaro and Rothwell, 2014; Lefaucheur et al., 2014; Tremblay et al., 2019). However, defining optimal targets for TMS remains a challenging task, which is mostly done based on brain’s morphology, using coordinates from atlases, or pre-defined locations from functional neuroimages (Cash et al., 2021).

With a stimulation coil placed on the scalp, TMS induces a brief electric field (E-field) that activates neurons in a limited brain region (approx. 1 cm^2^) (Terao and Ugawa, 2002). The localized activation, however, propagates to different areas via the white-matter network (Van Essen, 2013). Knowing which networks or connections of the brain are affected by TMS is important because structural connectivity of the brain plays a role in understanding and treating many brain disorders including MDD (Korgaonkar et al., 2014), Alzheimer’s disease (Lo et al., 2010), multiple sclerosis (Llufriu et al., 2017) and stroke (Yamada et al., 2004). Connections in the brain’s white matter can be detected non-invasively and in vivo with fiber tracking, *i.e.*, tractography, using diffusion magnetic resonance imaging (dMRI) (Shi and Toga, 2017). During the last years, research based on tractography has contributed substantially to elucidating the circuitry of the human brain (Wandell, 2016). Tractography can be performed on the whole brain, providing the structural connectome to study its network properties (Rubinov and Sporns, 2010), or on selected, *i.e.*, seed, regions in the brain.

State-of-the-art nTMS systems provide real-time updates of the coil position and the estimated induced E-field overlaid on the individual’s structural T1-weighted MRI (Ruohonen and Karhu, 2010; Hannula and Ilmoniemi, 2017). The addition of real-time tractography information to the already existing nTMS would be highly valuable by making it possible to aim for fiber tracts or to target regions that are remotely connected to the area under the coil. However, introducing real-time structural connectivity estimation to existing nTMS systems is challenging. This is mainly due to the inherent limitations involved with the accuracy of tractography, which have been increasingly pointed out in recent years by validation studies (Thomas et al., 2014; Yendiki et al., 2022) and benchmarks conducted through international tractography challenges (Maier-Hein et al., 2017; Nath et al., 2020; Schilling et al., 2021; Maffei et al., 2022). Importantly, tractography is well-known to miss connections that are present in the brain, *i.e.*, false negatives (Aydogan et al., 2018), and at the same time, it generates connections that do not exist, *i.e.*, false positives (Schilling et al., 2019).

The goal of this work was to develop a system that computes and displays brain’s structural connections in real time for guiding TMS. Our approach separates the slow, offline dMRI data pre-processing, and in real time, it computes the streamlines by our custom tractography algorithm (Aydogan and Shi, 2021) and displays the connections using an in-house developed neuronavigation system (Souza et al., 2018). By incorporating anatomical constraints, the proposed technique removes implausible streamlines, decreasing the output of false positive connections. Our visualization method also features a transfer function for streamline opacity, providing a visual feedback on the reliability of the displayed streamlines. In this article, we demonstrate the feasibility and real-time performance of our technique for TMS applications with synthetic data and in experiments with four healthy subjects.

## 2. Method

As shown in Fig. 1, our workflow is divided into two parts: an offline analysis (preparatory) and an online tractography computation with neuronavigation (real-time operation).

**Figure 1:**
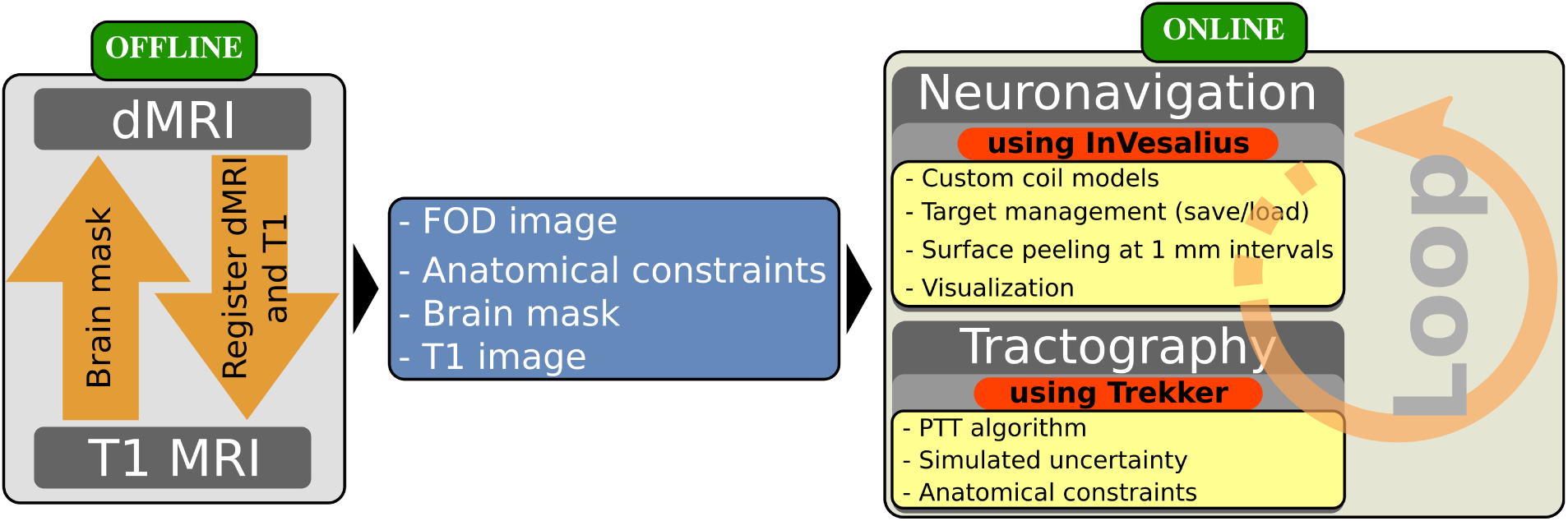
For real-time tractography-assisted neuronavigation, we combine information from T1 and dMRI data. In the offline (preparatory) part, necessary inputs for the online (real-time) part are prepared. The necessary inputs are: *i)* fiber orientation distribution (FOD) image, needed for fiber tracking, *ii)* anatomical constraints, needed to reduce false positives during tractography, *iii)* segmented brain mask, needed to compute the peeled brain surfaces, and *iv)* T1 image, to show the grayscale brain image on the peeled surfaces. The online part consists of the neuronavigation and tractography modules that continuously run in a multi-threaded loop. Neuronavigation and tractography are done using our custom software InVesalius (Souza et al., 2018) and Trekker (https://dmritrekker.github.io/), respectively. The main features used during real-time operation are inside the yellow boxes.

### 2.1. Offline processing

dMRI was denoised (Veraart et al., 2016) and corrected for artefacts induced by eddy currents and motion (Andersson and Sotiropoulos, 2016). Fiber orientation distribution (FOD) image was computed using the compartment model approach in Tran and Shi (2015). For anatomically constrained tractography (Smith et al., 2012), labels for white matter, cerebrospinal fluid (CSF), and the region outside the brain were defined from Freesurfer’s *reconall* (Fischl, 2012).

### 2.2. Online processing

#### 2.2.1. Tractography

##### Fiber tracking

Our parallel transport tractography (PTT) algorithm (Aydogan and Shi, 2019, 2021) is implemented in our in-house developed, open-source software Trekker (https://dmritrekker.github.io/). To reduce false positive connections, the following anatomical constraints are applied as *pathway rules* in Trekker:

1. To prevent improper termination *→ discard if ends inside* white matter
2. To prevent projecting through CSF *→ discard if enters* CSF
3. To prevent leaking outside the brain *→ stop at entry* outside the brain

##### Visualization of uncertainty

In this work, we introduce a novel visualization approach using the observations in Aydogan et al. (2018), which reports performance trends in tractography based on parameter combination choices. For example, a low FOD threshold parameter helps to find intricate connections, reducing false negatives, but at the same time this increases false positive streamlines. This prior information regarding the trade-off between sensitivity and specificity offers an opportunity to visualize uncertainty based on parameter choices. To visualize uncertainty in the fiber tracking results, we designed a transfer function that assigns each streamline an opacity value based on the fiber tracking parameter used to obtain that streamline. A precursory version of this approach was presented in ISMRM 2020 (Aydogan, 2020). Graphical explanation of this approach and the transfer function are shown in Fig. 2.

**Figure 2:**
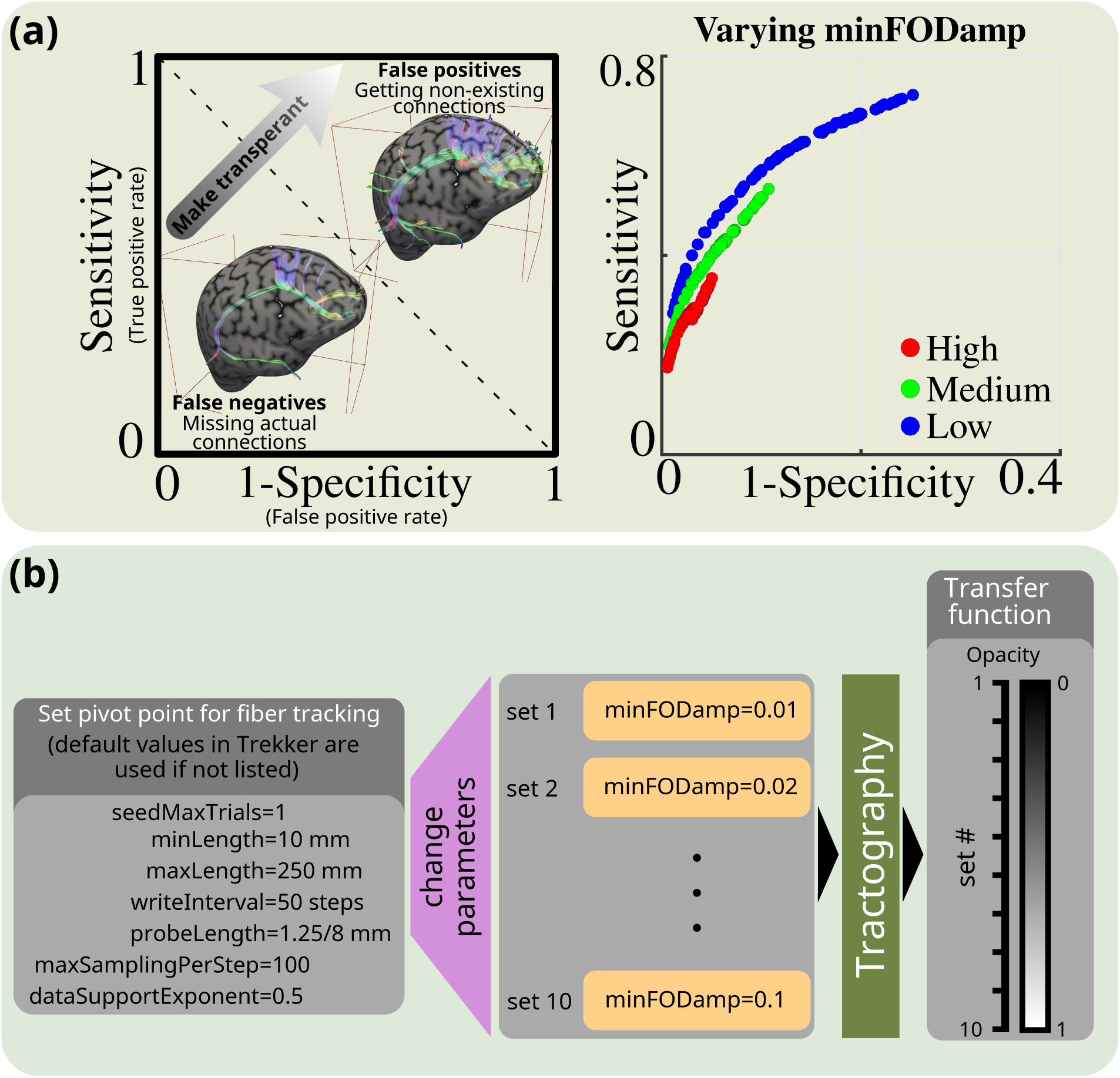
(a) Sensitivity and specificity plane to demonstrate the trade-off in tractography performance. The ROC curve can be traversed by varying the FOD threshold (*minFODamp* parameter in Trekker). Low values for FOD threshold lead to increased sensitivity at the cost of decreased specificity (Aydogan and Shi, 2018). By displaying the streamlines generated using low FOD thresholds with more transparency, we provide visual information to the operator regarding increased possibility of false connectivity as a result of the corresponding parameter choice. (b) Fixed parameters used for fiber tracking and the range of values for the varying FOD threshold as well as the corresponding transfer function, *i.e.*, the opacity used for the visualization.

#### 2.2.2. Implementation of real-time tractography during neuronavigation

The real-time visualization of tractograms during neuronavigation was achieved by integrating the Python API of Trekker in the open-source, free neuronavigation software InVesalius Navigator (Souza et al., 2018) (https://invesalius.github.io/). Tracking parameters are saved in a .json file and can be updated, to instantly fine-tune fiber tracking parameters. During neuronavigation, a continuous loop starts as soon as the TMS coil is detected by the tracking camera. Then, *N* seed coordinates for computing the streamlines are pseudo-randomly sampled from a sphere with 1.5-mm radius. The center of the sphere is defined as the closest white-matter point inside a rectangular prism created with its longitudinal axis coincident to a line projected from the TMS coil center. The rectangular prism has a square profile of 2*×*2 mm^2^, a height of 20 mm with grid points spaced by 1 mm, and is offset by 30 mm from the TMS coil surface. The *N* seed coordinates initiate the computation of *N* streamlines in each iteration; where *N* is automatically set as the number of computer’s processor’s threads. At each iteration, the *minFODamp* parameter is adjusted as shown in Fig. 2 and streamlines are visualized every 100 ms with the corresponding opacity. Because 10 parameter combinations are used for uncertainty visualization, these combinations are repeated at every 10^th^ iteration. The loop runs continuously until 1000 streamlines are displayed or the coil is moved by at least 2 mm. This distance threshold was implemented to avoid excessive removal and addition of streamlines for small jitters in the TMS coil coordinates. If the TMS coil moves more than 2 mm from the first point that the white matter was detected, streamlines that were previously displayed are removed, otherwise they are continuously added.

## 3. Experimental setup

### 3.1. Synthetic characterization

We studied the tracking parameters for uncertainty visualization with offline experiments conducted on the ISMRM 2015 tractography challenge dataset (Maier-Hein et al., 2017). The details about the data can be obtained from http://www.tractometer.org/ismrm_ 2015_challenge/data. We used the fiber tracking parameters shown in Fig. 2 to compute 10 million streamlines in the whole brain by randomly seeding the white-matter mask. The process is repeated for each of the 10 *minFODamp* values. Bundle overreach and overlap were compared against the submissions made to the original challenge.

### 3.2. TMS experiment

Experiments were done on four healthy male volunteers (age: 30–42). Written informed consents were collected from all participants. The study was done in accordance with the Declaration of Helsinki and approved by the Coordinating Ethics Committee of the Hospital District of Helsinki and Uusimaa.

#### 3.2.1. MRI data

MRI data were acquired using a MAGNETOM Skyra 3T MR scanner (Siemens Healthcare, Erlangen, Germany) with a 32-channel head coil. MRI measurements were done at the Advanced Magnetic Imaging Centre of Aalto NeuroImaging. For T1 image, a sagittal MPRAGE protocol with TE=3.3 ms, TR=2530 ms, 1*×*1*×*1 mm^3^ voxel dimension and 176*×*256*×*256 voxels were used. The dMRI were acquired according to a multi-shell high-angular resolution diffusion imaging (HARDI) scheme with TE=107 ms, TR=3.9 s, 2*×*2*×*2 mm^3^ voxel dimension and 176*×*256*×*256 voxels. Data were collected from a total of 100 gradient directions that are uniformly distributed on a sphere. 18, 32 and 50 volumes were distributed to three shells with *b*-values 900, 1600 and 2500 s/mm^2^, respectively. 11 *b*0 images were interleaved between the volumes and additional four *b*0 images were collected at the end according to a reverse phase-encoding scheme, for motion and distortion correction.

#### 3.2.2. TMS experimental protocol

Fig. 3 shows our setup. Navigation was performed with an infrared Polaris Vicra camera (Northern Digital Inc., Waterloo, ON, Canada) and tracking probes with passive reflective spherical markers. Fiducial registration errors were kept below 3 mm (Souza et al., 2018). Experiments were performed with a Dell Precision 7530 (CPU Intel 6 core 2.6 GHz i7-8850H, 32 GB RAM, 1TB SSD hard drive, NVidia Quadro P2000 graphics card, and Windows 10 64 bits).

**Figure 3:**
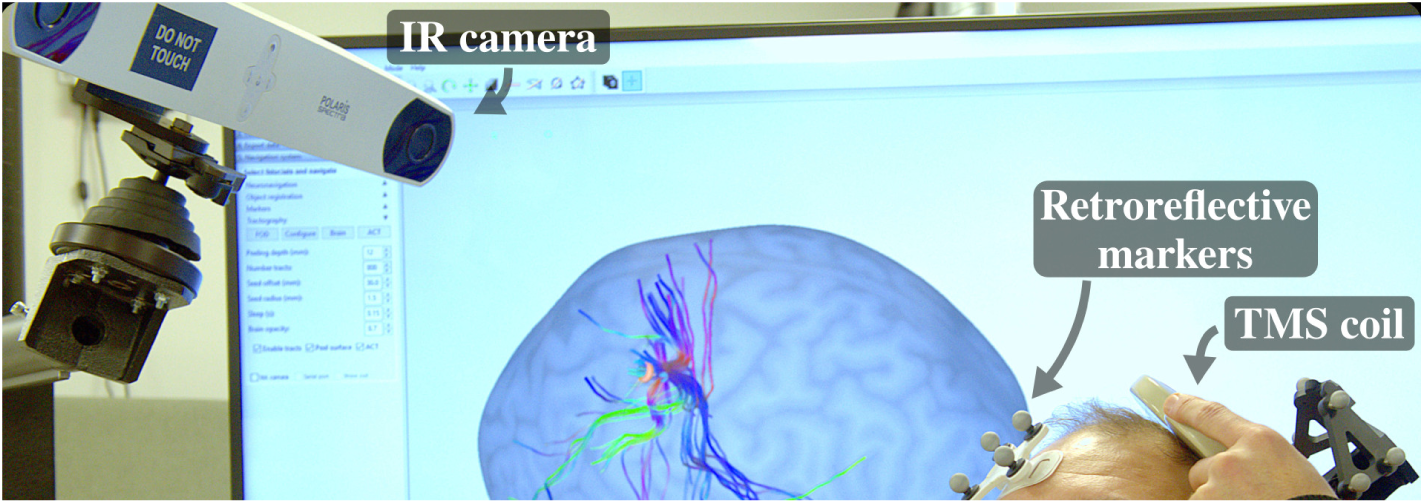
Tractography-assisted nTMS setup. The display with the InVesalius user interface shows the peeled brain surface and the streamlines obtained from the real-time TMS coil position. The coil is being held by the operator and tracked by the neuronavigation software that analyzes data from an infrared (IR) camera and retroreflective markers.

We studied the feasibility of reliable and repeatable targeting of four major connections in the left part of the brain involved in: motor, cognitive, speech, and visual functions. Therefore, we targeted the *i)* primary motor cortex (M1), *ii*) dorsolateral prefrontal cortex (DLPFC), *iii*) Broca’s area (BA44), and *iv*) primary visual cortex (V1), respectively.

We initially saved the intended TMS target locations related to these four regions in the InVesalius software interface to guide the TMS coil placement. The M1 target was identified based on abductor pollicis brevis (APB) muscle twitches. DLPFC was identified visually from T1 MRIs as described in Lioumis et al. (2009). Broca’s area was identified as described in speech cortical mapping studies (Lioumis et al., 2012; Corina et al., 2010; Krieg et al., 2017). The primary visual cortex was selected based on visual inspection of the anatomical MRIs and by applying single TMS pulses that elicited phosphenes in the participant’s visual field.

For assessing the reproducibility of the displayed streamlines, we placed the TMS coil on the selected target and recorded the TMS coil coordinates and corresponding seed coordinates used to compute the streamlines in real time. This process was repeated ten times for each of the four targets by removing the coil from the vicinity of participants’s head after each trial.

#### 3.2.3. Data analysis

The seed coordinates obtained during the experiments were used to study: **1)** how offline filtering (selection) of streamlines compare against real-time tractography, and **2)** the number of streamlines to display.

For **1)**, we generated 10 million streamlines for each of the 10 *minFODamp* values by randomly seeding the whole brain. Each tractogram was then filtered so that only those streamlines that passed through the 1.5-mm sphere centered at the recorded target points remained. For each target region, we reported the number of selected streamlines.

For **2)**, we investigated the overlap between the tractograms with increasing number of streamlines. For that, we first generated 100000 streamlines from each seed using each of the 10 *minFODamp* values. The combined tractogram with 1 million streamlines was used as a reference. We then simulated the case during the real-time experiment, where only a subset of these streamlines were shown. To that end, we generated 7 subsets with 100, 300, 1000, 3000, 10000, 30000 and 100000 streamlines. Each subset contained an equal number of streamlines computed with different *minFODamp* values. The overlap was computed by finding the intersection between thresholded (>0) track-density images (TDI) (Calamante et al., 2010).

## 4. Results

### 4.1. Synthetic characterization

Fig. 4a shows the overlap and overreach values with respect to *minFODamp*. The overlap shows how much the tractogram aligns with the ground truth, which is a measure of true positive connection. The overreach shows how much of the tractogram is outside the ground truth, which is a measure of false positive connections. In Fig. 4b, we picked three tractograms and combined them with and without uncertainty visualization. The three tractograms were obtained from whole-brain tractograms computed with *minFODamp* values of 0.01, 0.05 and 0.1, by selecting 500 streamlines within a sphere of radius 1.5-mm that was manually placed in the primary motor cortex.

**Figure 4:**
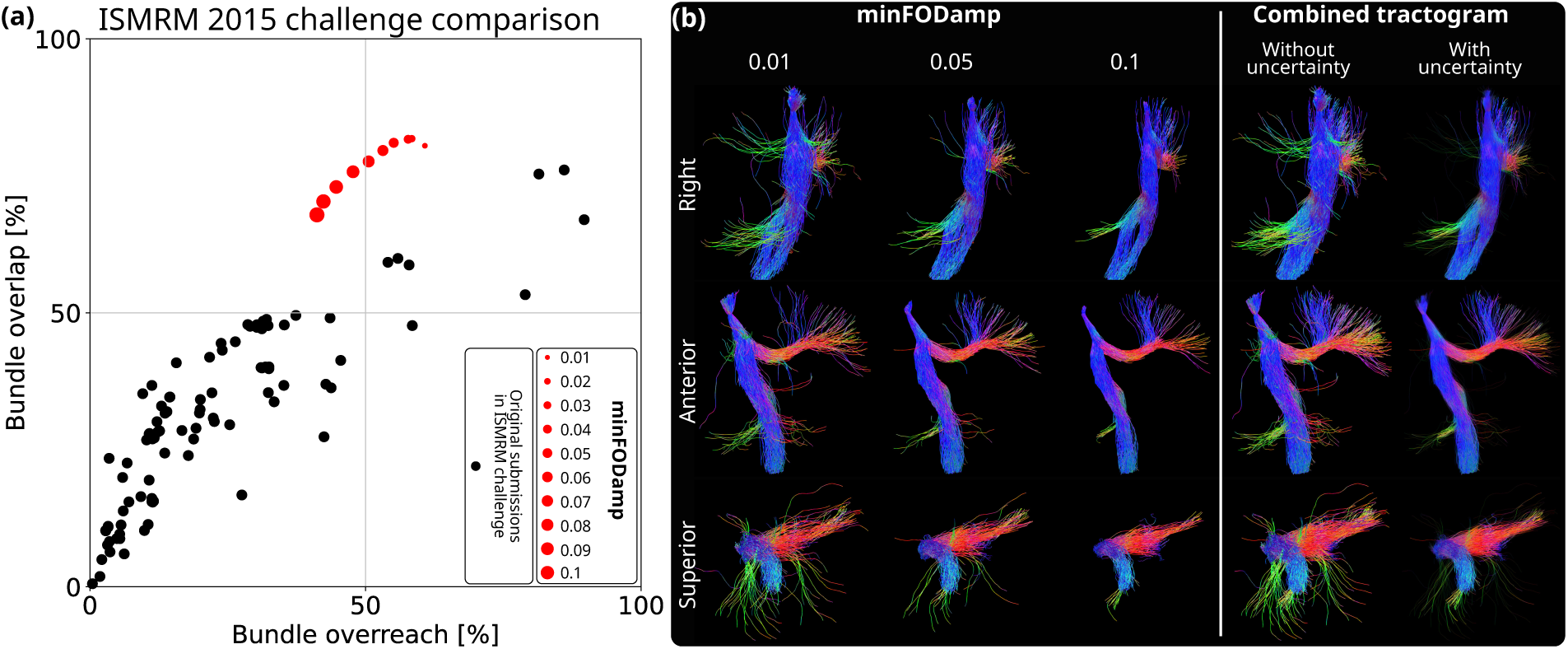
The variation in tractography performance for uncertainty visualization was tested using the ISMRM 2015 tractography challenge data. (a) Each red dot represents a score obtained for a tractogram with 10 million streamlines generated by randomly seeding the whole-brain mask using each of the 10 *minFODamp* parameters shown in Fig. 2. The obtained scores with Trekker are highly competitive against the original submissions to the challenge. The trend shows the expected trade-off between true and false positives, where increasing the *minFODamp* parameter decreases bundle overreach, *i.e.*, false positives, at the cost of reduced bundle overlap, *i.e.*, true positives. (b) Tractograms show that increasing *minFODamp* produces streamlines that may not sample the whole extent of connections. Tractograms with lower *minFODamp* values reach more regions; however, streamlines lose organization, which can lead to increased false positives. Combination of the tractograms with uncertainty visualization shows all the streamlines. But because streamlines computed with lower *minFODamp* are shown with more transparency, user is provided with visual information that these connections are more likely to be false positives than other streamlines shown on display.

### 4.2. Navigated TMS with real-time tractography

Our setup is shown in the supplementary video. The operator can observe the structural connections while the coil is moved. For demonstrative purposes, the operator shows the connections on various locations. Consent of the model is obtained to publish his face in the video.

### 4.3. TMS experiment

We compared the number of streamlines and overlap percentages using tractograms obtained offline. Fig. 5a shows the number of streamlines that were obtained by filtering offline computed, large-scale whole-brain tractograms that contain 100 million streamlines for each subject. We observe that there is large variability among subjects, brain regions, and repetitions. Nearly 15000 streamlines were obtained for many repetitions in the V1 area of subject #4. However, for the same area of subject #3, many times it was not possible to obtain any streamline. We observe a general trend of decreasing number of streamlines with the increase in *minFODamp*. For a few seed points, however, the number of streamlines increase with *minFODamp*, *e.g.*, M1 seed #4 of subject #2.

**Figure 5:**
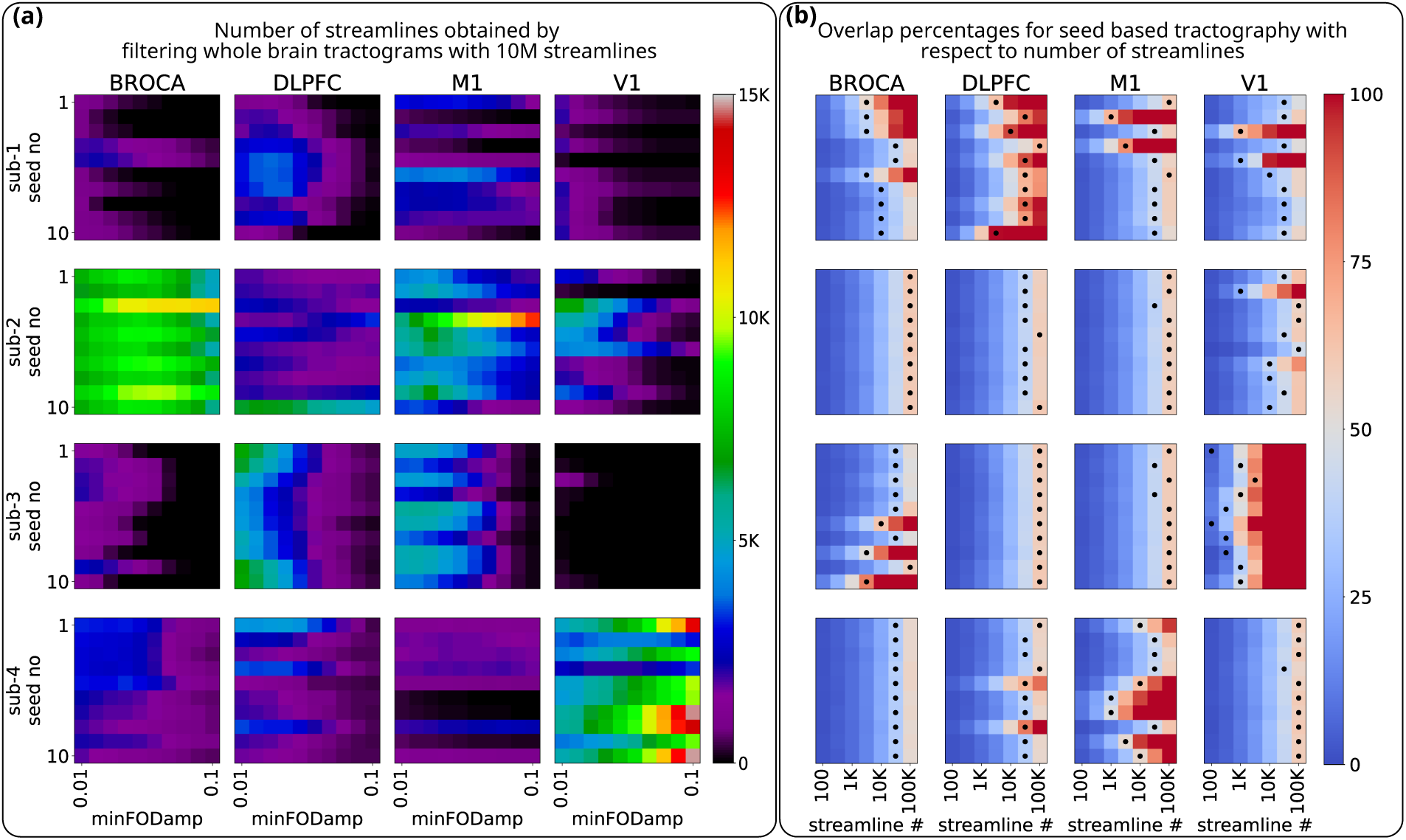
(a) shows the number of offline filtered streamlines. Each value is obtained by selecting streamlines that pass through the seed regions from a whole-brain tractogram containing 10 million streamlines. Values are reported for each of the 10 repetitions of coil placement, *i.e.*, seed point, as well as 10 different *minFODamp* values. (b) shows overlap percentages with respect to increasing the number of streamlines. Here, the streamlines are obtained offline with the same method as the real-time case. The reference contains 1 million streamlines obtained by combining 10 seed-based tractograms (see Section 3.2.3). Each of the 10 tractograms are computed with a different *minFODamp* value and contains 100000 streamlines. Black dots show the overlap obtained by filtering the whole-brain tractograms that contain 100 million streamlines for each seed.

In Fig. 5b, we showed the overlap values for the seed-based approach that is used for real-time tractography-based neuronavigation. The overlap percentages for increasing number of streamlines were computed for each seed, against the corresponding reference tractogram with 1 million streamlines (see Section 3.2.3). As expected, the overlap increases with the number of streamlines. For comparison, we also computed the overlap between the results of the offline filtering approach and the reference tractograms. Black dots in the figure show the closest overlap values obtained with the offline filtering results. The maximum number of dots is 64 at 30000 streamlines. There are 57 dots at 100000 thousand streamlines. Seed-based tractography covers a large portion of the target area after a few thousands of streamlines. This can be reached in a few seconds during real-time operation. When compared to the filtering approach, seed-based tractography covers a larger portion of the brain after a few tens of thousands of streamlines for many seed points, *e.g.*, Broca’s area of subject #1 or M1 of subject #4.

## 5. Discussion

We developed a real-time platform to compute and visualize structural connections in the brain with features tailored for guiding TMS applications. Our method uses state-of-the-art tractography practises that have received high scores in international challenges, yielding visual representations that are as accurate as possible. These practises include the use of: *i)* a modern FOD computation method that can handle complex white-matter fiber organizations (Tran and Shi, 2015), *ii)* application of anatomical constraints (Smith et al., 2012), and *iii)* a state-of-the-art fiber tracking algorithm (Aydogan and Shi, 2021). These practices provide improvements in the quality of tractograms compared to previous real-time tractography tools (Golby et al., 2011; Elhawary et al., 2011; Chamberland et al., 2014).

### Technical aspects of real-time tractography

There are two main ways to obtain the streamlines for visualization in real time. The first is to perform the fiber tracking in real time, as we did in our work. The second is to compute a large number of streamlines, and then select the relevant ones in real time. The ability to adjust fiber tracking parameters is arguably the most important benefit of our system. Because even for the same brain, optimal tractography parameters vary depending on the region or the white-matter tract (Aydogan et al., 2018). On the other hand, the availability of a pre-computed whole-brain tractogram enables to compute connectivity strengths and matrices (Daducci et al., 2016) — not possible to compute with real-time seed-based tractography.

Tractography algorithms are more complicated than filtering tractograms. However, real-time tractography has advantages when managing computer resources, because it does not require an interface between the hard drive and memory. On the other hand, large pre-computed whole-brain tractograms (with 100 million streamlines), can require 100 GB or more hard drive space, that can introduce challenges in clinics when transferring the data. Overall, both real-time tracking and filtering are suitable for tractography-assisted neuronavigation, and depending on the location in the brain, up to several hundreds or a few thousands streamlines per second can be obtained using standard computers. When compared against alternative offline practices, our real-time tractography pipeline does not compromise from quality in order to speed up the computation. Therefore, doing tractography in advance would not have improved the results that we show to the operator. Future research may lead to alternative visualization techniques that are capable of combining information from offline whole-brain tractograms to highlight higher-order connections of the seed region and its connectivity strength with the rest of the brain.

### Design aspects of streamline visualization and TMS neuronavigation

Our work distinguishes from previous works through the use of novel visualization techniques: *i)* Displaying tractograms with the peeled brain surface is a natural choice for TMS neuronavigation that was not demonstrated before. *ii)* Our dynamic, incremental, visualization of streamlines, distributes the complicated connectivity information along time, helping the operator to interpret the complex information. In contrast to previous visualization approaches that show a snapshot for connectivity (Golby et al., 2011; Elhawary et al., 2011; Chamberland et al., 2014), we are showing a movie, where thousands of streamlines can be displayed in an iterative and sequential fashion. *iii)* The uncertainty visualization approach is primarily designed for improved real-time experience, which to our knowledge, is the first time that transparency is used to convey information about uncertainty. For that, we first showed this trend using the ISMRM 2015 challenge data. While the exact overlap and overreach values shown in Fig. 4 are going to be different for other data, the performance trend is expected to be similar (Aydogan et al., 2018). As a result, our proposed uncertainty visualization provides a new insight about the reliability of the streamlines. While the current study develops new methods for visualization and provides a qualitative evaluation, future research should seek to answer whether the approaches quantitatively benefit the TMS operator during neuronavigation, for example, by improving treatment outcomes through individualized targeting.

### Impact of the seeding strategy

Our current setup estimates the seed region using the coil position and the white-matter segmentation. The brain areas affected by TMS can be better estimated with the E-field distribution (Weise et al., 2020; Aberra et al., 2020; Sollmann et al., 2016). Even so, the response to TMS is still uncertain, and the fiber orientations can play a role (Laakso et al., 2013). Therefore, the integration of real-time E-field estimates and tractography might improve the accuracy of future TMS-targeting methods.

Because TMS may primarily affect regions that are close to the coil (Siebner et al., 2022), gyral bias becomes a major problem for TMS neuronavigation with tractography. We believe this is reflected in the results shown in Fig. 5a. Even with a large number of streamlines (100 million), we observe that some regions were not reached by tractography, *e.g.*, V1 of subject #3. In Fig. 5, we not only observe poor streamline counts for some seeds, but we also see a large variability in the count that reach the seed regions for different subjects. For instance, while several thousands of streamlines could be obtained for most of the seeds in the Broca’s region of subject #2, no streamlines could be obtained for subject #1 and subject #3 for many seeds in the same region. Some of this variability may be due to differences in brain structure between individuals; however, we believe that the poor dMRI signal and fiber configuration variability around the cortex can be more significant factors. These highlight that even though we carefully adapted the state-of-the-art practices in our pipeline, there is room for improvement.

### Immediate applications of real-time tractography-assisted TMS neuronavigation

nTMS has been used with tractography to improve surgical outcomes (Picht et al., 2016) by identifying and visualizing eloquent motor areas during pre-operative planning (Frey et al., 2014), . This is achieved by finding and saving nTMS-based seed points in a disk or hospital’s picture archiving and communication system (PACS) (Mäkelä et al., 2015), followed by neurosurgeons’ using a separate software for seed-based tractography. Real-time tractography-assisted TMS neuronavigation can save time and costs by eliminating the need for a separate tractography step.

Our real-time tractography-assisted neuronavigation could be highly useful for paired associative-stimulation (PAS) (Koch and Rothwell, 2009; Koch et al., 2010). PAS has been shown to induce plastic changes (Classen et al., 2004), by involving stimulation of multiple targets, *e.g.*, two brain regions connected with cortico-cortical projections. Traditionally, one of the targets is set during the experiment based on functional measurements while the other targets are set manually based on anatomical MRI (Koch et al., 2013). Recently, Hernandez-Pavon et al. (2022) used dMRI-based tractography to post-hoc demonstrate that their stimulation sites were connected. Real-time tractography-assisted neuronavigation enables more precise and personalized PAS protocols in which connected sites can be identified during the experiment.

Recent developments in multi-channel TMS technology (Souza et al., 2022; Nieminen et al., 2022) opened a possibility for automated targeting (Tervo et al., 2022) and fast mapping of brain functions. Real-time tractography can play an important role for automated scanning algorithms to optimize stimulation parameters based on the underlying brain network. This would enable precise targeting of local and whole-brain networks for personalized connectomic neuromodulation (Horn and Fox, 2020).

### Limitations of tractography

We believe that the limited accuracy of tractography is the main challenge for the adaptation of real-time tractography-assisted neuronavigation. There are several factors that can negatively impact the reliability of tractograms. dMRI data acquired with low resolution and/or few diffusion samples (Calabrese et al., 2014), and sub-optimal pre-processing choices could lead to worse tractograms (Irfanoglu et al., 2012). It was shown that modern white-matter microstructure models, *e.g.*, those that can distinguish crossing fiber configurations, provide superior tractography results when compared against traditional techniques, such as the diffusion tensor imaging (DTI) (Farquharson et al., 2013). The choice of tractography algorithms (Sarwar et al., 2019), and the use of anatomical constraints were also shown to affect the results (Smith et al., 2012). Tractography is known to perform worse where fibers cross (Jeurissen et al., 2013). Moreover, the two-year long (2019–2020) IronTract challenge (https://irontract.mgh. harvard.edu/) show that fiber configurations that go beyond crossing, *e.g.*, fanning, branching, can be more challenging for tractography (Maffei et al., 2020, 2022; Schilling et al., 2022). Due to differences in fiber configuration in gray and white matters, and the folded geometry of the brain, tractography algorithms also tend to be biased towards terminating the streamlines at gyral crowns (Reveley et al., 2015). Overall, tractograms contain large amounts of false positives and false negatives that have been shown in several previous validation studies and tractography competitions (Thomas et al., 2014; Maier-Hein et al., 2017; Schilling et al., 2019; Aydogan et al., 2018; Girard et al., 2020; Maffei et al., 2020, 2022). While the aforementioned limitations impact the quality of tractography, we believe that our techniques represent a significant progress in tractography-assisted neuronavigation, proposing a solution that shows the brain’s anatomical connections in a way that is most accurate and helpful for brain stimulation, especially for TMS applications.

## 6. Conclusion

We developed a real-time tractography-assisted neuronavigation system for TMS. We anticipate that this technology is a critical step towards personalized brain stimulation targeting based on anatomical networks with potential applications in research and clinical environments.

## CRediT authorship contribution statement

**Dogu Baran Aydogan:** Conceptualization, Methodology, Software, Formal analysis, Investigation, Resources, Visualization, Writing - Original Draft, Writing – review & editing. **Victor H. Souza:** Conceptualization, Methodology, Software, Formal analysis, Investigation, Resources, Visualization, Writing - Original Draft, Writing – review & editing. **Renan H. Matsuda:** Software, Investigation, Writing - Review & Editing. **Pantelis Lioumis:** Conceptualization, Investigation, Resources, Writing - Original Draft, Writing – review & editing. **Risto J. Ilmoniemi:** Conceptualization, Resources, Writing - Review & Editing, Supervision, Project administration, Funding acquisition.

## Acknowledgements

This project has received funding from the Academy of Finland (grants #348631, #349985, and #353798), from Helsinki University Central Hospital (VTR grant #TYH2022224), and from the European Research Council (ERC) under the European Union’s Horizon 2020 research and innovation programme (ConnectToBrain; grant agreement no. 810377). RHM has received funding from the Conselho Nacional de Desenvolvimento Científico e Tecnoĺogico (grant no. 141056/2018-5) and from the FAPESP Research, Innovation and Dissemination Center for Neuromathematics (grant no. 2013/07699-0). Aalto NeuroImaging (ANI) infrastructure was used for MRI data collection at AMI, and for experiments at Aalto TMS. We thank Joonas Laurinoja and Tuomas Mutanen for their help with the video production. The authors acknowledge the computational resources provided by the Aalto Science-IT project.

## Notes

### Competing Interest Statement

PL has been consulting Nexstim Plc in matters other than diffusion based navigated TMS. RI has consulted Nexstim Plc and has several patents or patent applications related to TMS.

## References

Aberra, A.S., Wang, B., Grill, W.M., Peterchev, A.V., 2020. Simulation of transcranial magnetic stimulation in head model with morphologically-realistic cortical neurons. Brain Stimulation 13, 175–189. doi:https://doi.org/10.1016/j.brs.2019.10.002.

Andersson, J.L.R., Sotiropoulos, S.N., 2016. An integrated approach to correction for off-resonance effects and subject movement in diffusion MR imaging. NeuroImage 125, 1063–1078. doi:10.1016/j.neuroimage.2015.10.019.

Aydogan, D.B., 2020. Visualization of uncertainty in tractograms using ROC-based transfer functions for real-time TMS applications, in: Proceedings of the 28^th^ Annual Meeting of ISMRM. Online.

Aydogan, D.B., Jacobs, R., Dulawa, S., Thompson, S.L., Francois, M.C., Toga, A.W., Dong, H., Knowles, J.A., Shi, Y., 2018. When tractography meets tracer injections: a systematic study of trends and variation sources of diffusion-based connectivity. Brain Structure and Function 223, 2841–2858. doi:10.1007/s00429-018-1663-8.

Aydogan, D.B., Shi, Y., 2018. Tracking and validation techniques for topographically organized tractography. NeuroImage 181, 64–84. doi:10.1016/j.neuroimage.2018.06.071.

Aydogan, D.B., Shi, Y., 2019. A novel fiber-tracking algorithm using parallel transport frames, in: Proceedings of the 27^th^ Annual Meeting of ISMRM. Montreal.

Aydogan, D.B., Shi, Y., 2021. Parallel transport tractography. IEEE Transactions on Medical Imaging 40, 635–647. doi:10.1109/TMI.2020.3034038.

Calabrese, E., Badea, A., Coe, C.L., Lubach, G.R., Styner, M.A., Johnson, G.A., 2014. Investigating the tradeoffs between spatial resolution and diffusion sampling for brain mapping with diffusion tractography: Time well spent? Human Brain Mapping 35, 5667–5685.

Calamante, F., Tournier, J.D., Jackson, G.D., Connelly, A., 2010. Track-density imaging (TDI): Super-resolution white matter imaging using whole-brain track-density mapping. NeuroImage 53, 1233–1243.

Cash, R.F., Cocchi, L., Lv, J., Fitzgerald, P.B., Zalesky, A., 2021. Functional magnetic resonance imaging–guided personalization of transcranial magnetic stimulation treatment for depression. JAMA Psychiatry 78, 337–339. doi:10.1001/jamapsychiatry.2020.3794.

Chamberland, M., Whittingstall, K., Fortin, D., Mathieu, D., Descoteaux, M., 2014. Real-time multi-peak tractography for instantaneous connectivity display. Frontiers in Neuroinformatics 8, 59.

Classen, J., Wolters, A., Stefan, K., Wycislo, M., Sandbrink, F., Schmidt, A., Kunesch, E., 2004. Paired associative stimulation. Supplements to Clinical neurophysiology 57, 563–569.

Corina, D.P., Loudermilk, B.C., Detwiler, L., Martin, R.F., Brinkley, J.F., Ojemann, G., 2010. Analysis of naming errors during cortical stimulation mapping: implications for models of language representation. Brain and Language 115, 101–112.

Daducci, A., Dal Palú, A., Descoteaux, M., Thiran, J.P., 2016. Microstructure informed tractography: pitfalls and open challenges. Frontiers in Neuroscience 10, 247.

Di Lazzaro, V., Rothwell, J.C., 2014. Corticospinal activity evoked and modulated by non-invasive stimulation of the intact human motor cortex. The Journal of Physiology 592, 4115–4128. doi:https://doi.org/10.1113/jphysiol.2014.274316.

Elhawary, H., Liu, H., Patel, P., Norton, I., Rigolo, L., Papademetris, X., Hata, N., Golby, A.J., 2011. Intraoperative real-time querying of white matter tracts during frameless stereotactic neuronavigation. Neurosurgery 68, 506–516.

Farquharson, S., Tournier, J.D., Calamante, F., Fabinyi, G., Schneider-Kolsky, M., Jackson, G.D., Connelly, A., 2013. White matter fiber tractography: why we need to move beyond DTI. Journal of Neurosurgery 118, 1367–1377.

Fischl, B., 2012. Freesurfer. NeuroImage 62, 774–781.

Fitzgerald, P.B., Hoy, K., McQueen, S., Maller, J.J., Herring, S., Segrave, R., Bailey, M., Been, G., Kulkarni, J., Daskalakis, Z.J., 2009. A randomized trial of rTMS targeted with MRI based neuro-navigation in treatment-resistant depression. Neuropsychopharmacology 34, 1255–1262.

Frey, D., Schilt, S., Strack, V., Zdunczyk, A., Röosler, J., Niraula, B., Vajkoczy, P., Picht, T., 2014. Navigated transcranial magnetic stimulation improves the treatment outcome in patients with brain tumors in motor eloquent locations. Neuro-oncology 16, 1365– 1372.

Girard, G., Caminiti, R., Battaglia-Mayer, A., St-Onge, E., Ambrosen, K.S., Eskildsen, S.F., Krug, K., Dyrby, T.B., Descoteaux, M., Thiran, J.P., et al., 2020. On the cortical connectivity in the macaque brain: A comparison of diffusion tractography and histological tracing data. NeuroImage 221, 117201.

Golby, A.J., Kindlmann, G., Norton, I., Yarmarkovich, A., Pieper, S., Kikinis, R., 2011. Interactive diffusion tensor tractography visualization for neurosurgical planning. Neurosurgery 68, 496–505.

Grosprêtre, S., Ruffino, C., Lebon, F., 2016. Motor imagery and cortico-spinal excitability: a review. European Journal of Sport Science 16, 317–324.

Hannula, H., Ilmoniemi, R.J., 2017. Basic principles of navigated TMS. Springer International Publishing, Cham. pp. 3–29. doi:10.1007/978-3-319-54918-7_1.

Hernandez-Pavon, J.C., Schneider-Garces, N., Begnoche, J.P., Miller, L.E., Raij, T., 2022. Targeted modulation of human brain interregional effective connectivity with spike-timing dependent plasticity. Neuromodulation: Technology at the Neural Interface doi:https://doi.org/10.1016/j.neurom.2022.10.045.

Horn, A., Fox, M.D., 2020. Opportunities of connectomic neuromodulation. NeuroImage 221, 117180. doi:https://doi.org/10.1016/j.neuroimage.2020.117180.

Irfanoglu, M.O., Walker, L., Sarlls, J., Marenco, S., Pierpaoli, C., 2012. Effects of image distortions originating from susceptibility variations and concomitant fields on diffusion MRI tractography results. NeuroImage 61, 275–288.

Jeurissen, B., Leemans, A., Tournier, J.D., Jones, D.K., Sijbers, J., 2013. Investigating the prevalence of complex fiber configurations in white matter tissue with diffusion magnetic resonance imaging. Human Brain Mapping 34, 2747–2766.

Koch, G., Cercignani, M., Pecchioli, C., Versace, V., Oliveri, M., Caltagirone, C., Rothwell, J., Bozzali, M., 2010. In vivo definition of parieto-motor connections involved in planning of grasping movements. NeuroImage 51, 300–312.

Koch, G., Ponzo, V., Di Lorenzo, F., Caltagirone, C., Veniero, D., 2013. Hebbian and anti-hebbian spike-timing-dependent plasticity of human cortico-cortical connections. Journal of Neuroscience 33, 9725–9733.

Koch, G., Rothwell, J.C., 2009. TMS investigations into the task-dependent functional interplay between human posterior parietal and motor cortex. Behavioural Brain Research 202, 147–152.

Korgaonkar, M.S., Fornito, A., Williams, L.M., Grieve, S.M., 2014. Abnormal structural networks characterize major depressive disorder: a connectome analysis. Biological Psychiatry 76, 567–574.

Krieg, S.M., Lioumis, P., Mäkelä, J.P., Wilenius, J., Karhu, J., Hannula, H., Savolainen, P., Lucas, C.W., Seidel, K., Laakso, A., et al., 2017. Protocol for motor and language mapping by navigated TMS in patients and healthy volunteers; workshop report. Acta Neurochirurgica 159, 1187–1195.

Laakso, I., Hirata, A., Ugawa, Y., 2013. Effects of coil orientation on the electric field induced by TMS over the hand motor area. Physics in Medicine & Biology 59, 203.

Lefaucheur, J.P., André-Obadia, N., Antal, A., Ayache, S.S., Baeken, C., Benninger, D.H., Cantello, R.M., Cincotta, M., de Carvalho, M., De Ridder, D., et al., 2014. Evidence-based guidelines on the therapeutic use of repetitive transcranial magnetic stimulation (rTMS). Clinical Neurophysiology 125, 2150–2206.

Lefaucheur, J.P., Picht, T., 2016. The value of preoperative functional cortical mapping using navigated TMS. Neurophysiologie Clinique/Clinical Neurophysiology 46, 125–133.

Lioumis, P., Kičić, D., Savolainen, P., Mäkelä, J.P., Kähkönen, S., 2009. Reproducibility of TMS-evoked EEG responses. Human Brain Mapping 30, 1387–1396.

Lioumis, P., Zhdanov, A., Mäkelä, N., Lehtinen, H., Wilenius, J., Neuvonen, T., Hannula, H., Deletis, V., Picht, T., Mäkelä, J.P., 2012. A novel approach for documenting naming errors induced by navigated transcranial magnetic stimulation. Journal of Neuroscience Methods 204, 349–354.

Llufriu, S., Martinez-Heras, E., Solana, E., Sola-Valls, N., Sepulveda, M., Blanco, Y., Martinez-Lapiscina, E.H., Andorra, M., Villoslada, P., Prats-Galino, A., Saiz, A., 2017. Structural networks involved in attention and executive functions in multiple sclerosis. NeuroImage: Clinical 13, 288–296. doi:10.1016/j.nicl.2016.11.026.

Lo, C.Y., Wang, P.N., Chou, K.H., Wang, J., He, Y., Lin, C.P., 2010. Diffusion tensor tractography reveals abnormal topological organization in structural cortical networks in Alzheimer’s disease. Journal of Neuroscience 30, 16876–16885.

Maffei, C., Girard, G., Schilling, K.G., Adluru, N., Aydogan, D.B., Hamamci, A., Yeh, F.C., Mancini, M., Wu, Y., Sarica, A., et al., 2020. The IronTract challenge: Validation and optimal tractography methods for the HCP diffusion acquisition scheme. ISMRM, Virtual .

Maffei, C., Girard, G., Schilling, K.G., Aydogan, D.B., Adluru, N., Zhylka, A., Wu, Y., Mancini, M., Hamamci, A., Sarica, A., et al., 2022. Insights from the IronTract challenge: optimal methods for mapping brain pathways from multi-shell diffusion MRI. NeuroImage 257, 119327.

Maier-Hein, K.H., Neher, P.F., Houde, J.C., Côté, M.A., Garyfallidis, E., Zhong, J., Chamberland, M., Yeh, F.C., Lin, Y.C., Ji, Q., et al., 2017. The challenge of mapping the human connectome based on diffusion tractography. Nature Communications 8, 1–13. doi:10.1038/s41467-017-01285-x.

Mäkelä, T., Vitikainen, A.M., Laakso, A., Mäkelä, J.P., 2015. Integrating nTMS data into a radiology picture archiving system. Journal of Digital Imaging 28, 428–432. doi:10.1007/s10278-015-9768-6.

Nath, V., Schilling, K.G., Parvathaneni, P., Huo, Y., Blaber, J.A., Hainline, A.E., Barakovic, M., Romascano, D., Rafael-Patino, J., Frigo, M., et al., 2020. Tractography reproducibility challenge with empirical data (TraCED): The 2017 ISMRM diffusion study group challenge. Journal of Magnetic Resonance Imaging 51, 234–249.

Nieminen, J.O., Sinisalo, H., Souza, V.H., Malmi, M., Yuryev, M., Tervo, A.E., Stenroos, M., Milardovich, D., Korhonen, J.T., Koponen, L.M., Ilmoniemi, R.J., 2022. Multi-locus transcranial magnetic stimulation system for electronically targeted brain stimulation. Brain Stimulation 15, 116–124. doi:10.1016/j.brs.2021.11.014.

Pelissolo, A., Harika-Germaneau, G., Rachid, F., Gaudeau-Bosma, C., Tanguy, M.L., BenAdhira, R., Bouaziz, N., Popa, T., Wassouf, I., Saba, G., et al., 2016. Repetitive transcranial magnetic stimulation to supplementary motor area in refractory obsessive-compulsive disorder treatment: a sham-controlled trial. International Journal of Neuropsychopharmacology 19, pyw025.

Picht, T., Frey, D., Thieme, S., Kliesch, S., Vajkoczy, P., 2016. Presurgical navigated TMS motor cortex mapping improves outcome in glioblastoma surgery: a controlled observational study. Journal of Neuro-Oncology 126, 535–543. doi:10.1007/s11060-015-1993-9.

Reveley, C., Seth, A.K., Pierpaoli, C., Silva, A.C., Yu, D., Saunders, R.C., Leopold, D.A., Ye, F.Q., 2015. Superficial white matter fiber systems impede detection of long-range cortical connections in diffusion MR tractography. Proceedings of the National Academy of Sciences 112, E2820–E2828.

Rubinov, M., Sporns, O., 2010. Complex network measures of brain connectivity: Uses and interpretations. NeuroImage 52, 1059–1069.

Ruohonen, J., Karhu, J., 2010. Navigated transcranial magnetic stimulation. Clinical Neurophysiology 40, 7–17.

Sarwar, T., Ramamohanarao, K., Zalesky, A., 2019. Mapping connectomes with diffusion MRI: deterministic or probabilistic tractography? Magnetic Resonance in Medicine 81, 1368–1384.

Schilling, K.G., Nath, V., Hansen, C., Parvathaneni, P., Blaber, J., Gao, Y., Neher, P., Aydogan, D.B., Shi, Y., Ocampo-Pineda, M., et al., 2019. Limits to anatomical accuracy of diffusion tractography using modern approaches. NeuroImage 185, 1–11. doi:10.1016/j.neuroimage.2018.10.029.

Schilling, K.G., Rheault, F., Petit, L., Hansen, C.B., Nath, V., Yeh, F.C., Girard, G., Barakovic, M., Rafael-Patino, J., Yu, T., et al., 2021. Tractography dissection variability: What happens when 42 groups dissect 14 white matter bundles on the same dataset? NeuroImage 243, 118502.

Schilling, K.G., Tax, C.M., Rheault, F., Landman, B.A., Anderson, A.W., Descoteaux, M., Petit, L., 2022. Prevalence of white matter pathways coming into a single white matter voxel orientation: The bottleneck issue in tractography. Human Brain Mapping 43, 1196–1213.

Shi, Y., Toga, A., 2017. Connectome imaging for mapping human brain pathways. Molecular Psychiatry 22, 1230–1240.

Siebner, H.R., Funke, K., Aberra, A.S., Antal, A., Bestmann, S., Chen, R., Classen, J., Davare, M., Di Lazzaro, V., Fox, P.T., et al., 2022. Transcranial magnetic stimulation of the brain: What is stimulated?–a consensus and critical position paper. Clinical Neurophysiology .

Smith, R.E., Tournier, J.D., Calamante, F., Connelly, A., 2012. Anatomically-constrained tractography: improved diffusion MRI streamlines tractography through effective use of anatomical information. NeuroImage 62, 1924–1938.

Sollmann, N., Goblirsch-Kolb, M.F., Ille, S., Butenschoen, V.M., Boeckh-Behrens, T., Meyer, B., Ringel, F., Krieg, S.M., 2016. Comparison between electric-field-navigated and line-navigated TMS for cortical motor mapping in patients with brain tumors. Acta Neurochirurgica 158, 2277–2289.

Souza, V.H., Matsuda, R.H., Peres, A.S., Amorim, P.H.J., Moraes, T.F., Silva, J.V.L., Baffa, O., 2018. Development and characterization of the InVesalius Navigator software for navigated transcranial magnetic stimulation. Journal of Neuroscience Methods 309, 109–120.

Souza, V.H., Nieminen, J.O., Tugin, S., Koponen, L.M., Baffa, O., Ilmoniemi, R.J., 2022. TMS with fast and accurate electronic control: Measuring the orientation sensitivity of corticomotor pathways. Brain Stimulation 15, 306–315. doi:https://doi.org/10.1016/j.brs.2022.01.009.

Terao, Y., Ugawa, Y., 2002. Basic mechanisms of TMS. Journal of Clinical Neurophysiology 19, 322.

Tervo, A.E., Nieminen, J.O., Lioumis, P., Metsomaa, J., Souza, V.H., Sinisalo, H., Stenroos, M., Sarvas, J., Ilmoniemi, R.J., 2022. Closed-loop optimization of transcranial magnetic stimulation with electroencephalography feedback. Brain Stimulation 15, 523– 531. doi:https://doi.org/10.1016/j.brs.2022.01.016.

Thomas, C., Ye, F.Q., Irfanoglu, M.O., Modi, P., Saleem, K.S., Leopold, D.A., Pierpaoli, C., 2014. Anatomical accuracy of brain connections derived from diffusion MRI tractography is inherently limited. Proceedings of the National Academy of Sciences 111, 16574–16579.

Tran, G., Shi, Y., 2015. Fiber orientation and compartment parameter estimation from multi-shell diffusion imaging. IEEE Transactions on Medical Imaging 34, 2320–2332.

Tremblay, S., Rogasch, N.C., Premoli, I., Blumberger, D.M., Casarotto, S., Chen, R., Di Lazzaro, V., Farzan, F., Ferrarelli, F., Fitzgerald, P.B., et al., 2019. Clinical utility and prospective of TMS-EEG. Clinical Neurophysiology 130, 802–844.

Van Essen, D.C., 2013. Cartography and connectomes. Neuron 80, 775–790. doi:10.1016/j.neuron.2013.10.027.

Veraart, J., Novikov, D.S., Christiaens, D., Ades-aron, B., Sijbers, J., Fieremans, E., 2016. Denoising of diffusion MRI using random matrix theory. NeuroImage 142, 394– 406. doi:10.1016/j.neuroimage.2016.08.016.

Wandell, B.A., 2016. Clarifying human white matter. Annual Review of Neuroscience 39, 103–128.

Weise, K., Numssen, O., Thielscher, A., Hartwigsen, G., Knösche, T.R., 2020. A novel approach to localize cortical tms effects. NeuroImage 209, 116486. doi:https://doi.org/10.1016/j.neuroimage.2019.116486.

Yamada, K., Ito, H., Nakamura, H., Kizu, O., Akada, W., Kubota, T., Goto, M., Konishi, J., Yoshikawa, K., Shiga, K., Nakagawa, M., Mori, S., Nishimura, T., 2004. Stroke patients’ evolving symptoms assessed by tractography. Journal of Magnetic Resonance Imaging 20, 923–929. doi:10.1002/jmri.20215.

Yendiki, A., Aggarwal, M., Axer, M., Howard, A.F., van Cappellen van Walsum, A.M., Haber, S.N., 2022. Post mortem mapping of connectional anatomy for the validation of diffusion MRI. NeuroImage 256, 119146. doi:https://doi.org/10.1016/j.neuroimage.2022.119146.

